# Circadian disruption enhances HSF1 signaling and tumorigenesis in Kras-driven lung cancer

**DOI:** 10.1101/2022.01.31.478213

**Authors:** Marie Pariollaud, Lara H. Ibrahim, Emanuel Irizarry, Rebecca M. Mello, Alanna B. Chan, Brian J. Altman, Reuben J. Shaw, Michael J. Bollong, R. Luke Wiseman, Katja A. Lamia

## Abstract

Disrupted circadian rhythmicity is a prominent feature of modern society and has been designated as a probable carcinogen by the World Health Organization. However, the biological mechanisms that connect circadian disruption and cancer risk remain largely undefined. We demonstrate that exposure to chronic circadian disruption (chronic jetlag, CJL) increases tumor burden in a mouse model of KRAS-driven lung cancer. Molecular characterization of tumors and tumor-bearing lung tissues revealed that CJL enhances the expression of heat shock factor 1 (HSF1) target genes. Consistently, exposure to CJL disrupted the highly rhythmic nuclear trafficking of HSF1 in the lung, resulting in an enhanced accumulation of HSF1 in the nucleus. HSF1 has been shown to promote tumorigenesis in other systems, and we find that pharmacological inhibition of HSF1 reduces the growth of KRAS-mutant human lung cancer cells. These findings implicate HSF1 as a molecular link between circadian disruption and enhanced tumorigenesis.

## Introduction

In 2015, the National Health Interview Survey revealed that 12–35% of the workforce in various United States industries work irregular schedules, including night and rotating shifts (*1*). Several human and animal studies have demonstrated that disruption of circadian rhythms, either by genetic or environmental means, enhances cancer risk (*2*), including the risk of lung adenocarcinoma (*3-7*). Lung cancer is the leading cause of cancer deaths in men and women worldwide (*8, 9*). The lung adenocarcinoma (LUAD) subtype of non-small cell lung cancer (NSCLC) is the most prevalent form of lung cancer and Kirsten rat sarcoma (KRAS) is the most frequently mutated oncogene in human LUAD (*10*). Despite extensive characterization of genetic events that contribute to lung cancer, there has been relatively little research addressing the impact of environmental circadian disruption on lung tumorigenesis in humans. The lung is under tight circadian control, as evidenced by robust 24-hour rhythms in intrinsic defense mechanisms and lung physiology indices such as lung resistance and peak expiratory flow (*11*). Selective ablation of bronchiolar epithelial cells results in the loss of circadian clock oscillations in mouse lung slices, demonstrating that airway epithelial cells are key circadian oscillators within the lung (*12*). NSCLC is a cancer of epithelial origin (*13, 14*), so this reinforces the hypothesis that disruption of the circadian machinery could trigger harmful events due to dysregulation of homeostasis, resulting in increased risk of lung tumor development.

The mammalian circadian machinery consists of an autoregulatory transcription-translation feedback loop. Its positive arm — heterodimers of circadian locomotor output cycles kaput (CLOCK) and brain and muscle ARNT-like protein 1 (BMAL1) — drives the transcription of two inhibitory arms — periods (PERs) and cryptochromes (CRYs) on one hand and nuclear receptor subfamily 1 group D members 1 and 2 (NR1D1/NR1D2 also called REV-ERBα/REV-ERBβ) on the other. While PERs and CRYs inhibit the BMAL1-CLOCK heterodimer transactivation function, REV-ERBs repress BMAL1 expression. These transcription-translation feedback loops drive 24-hour periodic expression of gene products leading to rhythmic physiologic functions. Accumulating evidence demonstrates that circadian clock components play critical roles in regulating several hallmarks of cancer, including control of cell proliferation, cell death, DNA repair, and metabolic alteration (*15-17*). However, the precise mechanisms underlying the cooperation between circadian clock disruption and tumorigenesis remain poorly understood. By manipulating lighting schedules to mimic the circadian disturbance that humans encounter during rotating shift work or frequent eastbound transmeridian flights (chronic jetlag, CJL), we show that this environmental light disruption alters gene expression in liver and lungs of mice. Further, we show that *Kras*^*LSL-G12D/+*^ (K) mice, a genetically engineered mouse model (GEMM) of NSCLC (*18*), developed many more tumors when housed in CJL compared to normal light conditions (12 hours of light; 12 hours of darkness; 12:12LD). Unbiased RNA sequencing and gene expression analyses revealed a profound disruption of the circadian clock machinery and chronic elevation of the heat shock response in lungs of mice exposed to CJL, indicating that light-induced circadian disruption perturbs homeostatic regulation of HSF1. Given the strong and growing evidence that HSF1 can facilitate tumorigenesis (*19-22*), these findings suggest that chronic elevation of HSF1 signaling could be a key molecular link between circadian disruption and increased cancer risk.

## Results

### Experimental Chronic Jetlag Disrupts Peripheral Clocks

To mimic chronic disruption of a functional circadian timing system, we used a chronic jetlag (CJL) protocol consisting of an 8-hour light phase advance repeated every 2 or 3 days (Fig. 1A). This altered light/dark scheme mimics the circadian disturbance that humans face during rotating shift work (*23, 24*) and has been shown to increase tumor development upon Glasgow osteosarcoma inoculation (*25*), in chemically-induced or spontaneous mouse liver cancer models (*26, 27*), and in a mouse model of lung cancer similar to the one used here (*3*). To first assess the magnitude of the disturbance of this protocol on the clock machinery and other cancer-related pathways, male and female C57BL/6J mice were housed in either normal light (12:12LD) or CJL conditions for 8 weeks before tissues were collected every 4 hours over a 24-hour period. Importantly, the timing of light-dark transitions on the day of collection (Day 1 in Fig. 1A) were the same for at least 24 hours before the collection (Day 7 in Fig. 1A). As expected, CJL greatly impacted the expression of core clock genes in lung, liver, and to a lesser extent spleen, with loss of periodic expression (P^JTKCycle^ < 0.05) for most of the genes, including *Bmal1, Cry1* and *Rev-Erba* (Fig. 1B, fig. S1). Similarly, the components of the clock machinery were profoundly disrupted at the protein level (Fig. 1C, fig. S1C). While these data are consistent with a model in which circadian rhythms of gene expression are suppressed or disrupted by CJL, we cannot exclude the possibility that the loss of observed rhythmicity for some genes is caused by a lack of synchrony between animals or an inability to properly assign individual animals to a specific circadian “phase” rather than loss of rhythmicity within individual mice based on this analysis. Nonetheless, the observation that *Per2* remains rhythmic in mice exposed to CJL while *Rev-Erb*α for example does not, suggests a desynchronization of the core clock components relative to each other. Interestingly, CJL exposure impacted daily rhythms of core clock genes somewhat differently in male versus female mice (fig. S1). These disparities are consistent with previous studies suggesting sexual dimorphism in circadian clock mechanism and physiology (*28*).

**Fig. 1.**
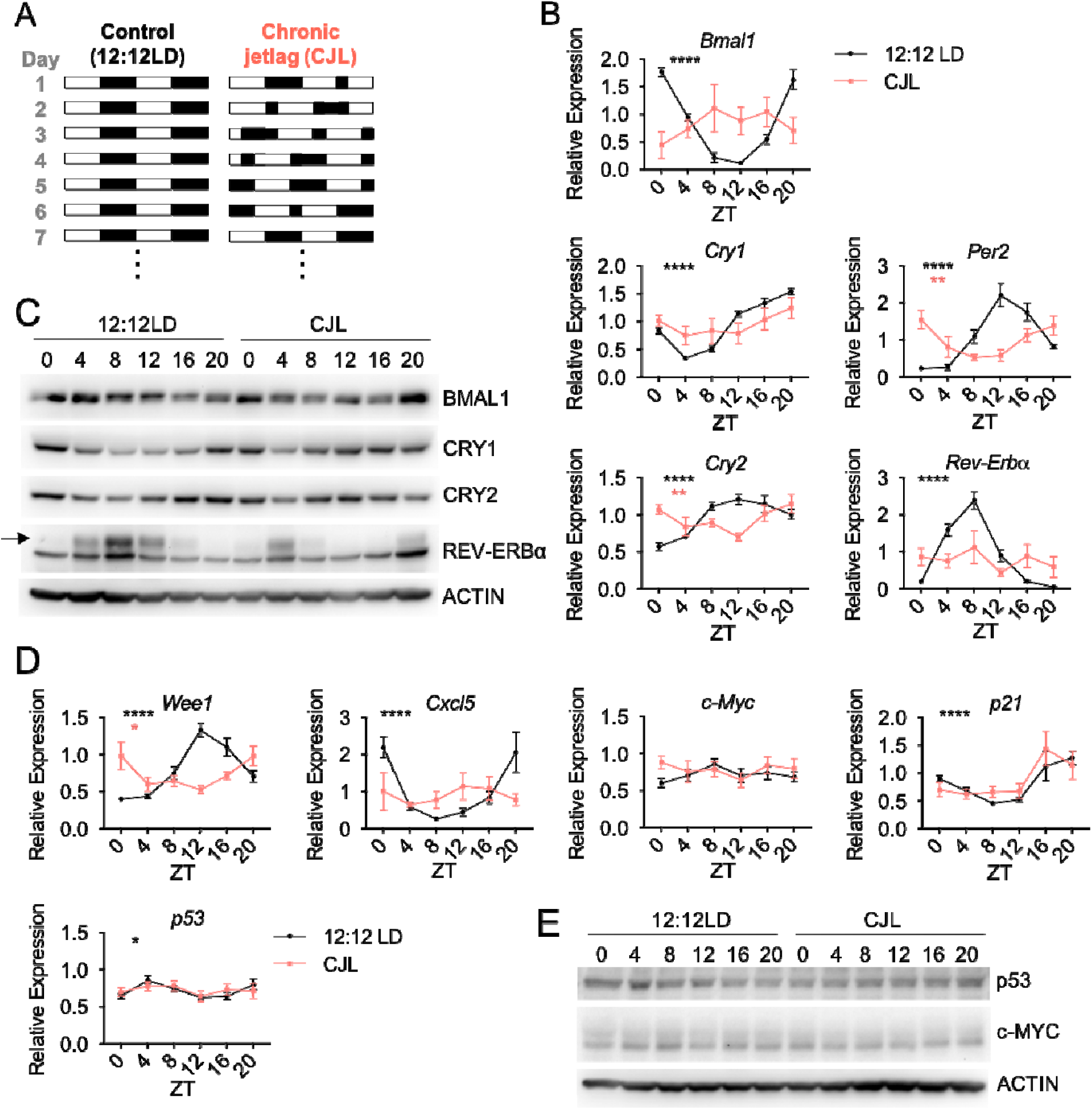
Chronic jetlag (CJL) severely impairs rhythmicity and magnitude of core clock and clock-controlled genes in the lung. **(A)** Schematic representation of the CJL protocol. White and black rectangles represent periods of light and dark, respectively. Each row represents two consecutive days starting with the numbered day shown at left. **(B-E**) C57BL/6J male and female mice were housed in 12:12LD or CJL for 8 weeks. Lung tissues were collected at the indicated times (ZT0: light on, ZT12: light off) on Day 1 of the schedule shown in (A). (**B**,**D**) Gene expression normalized to *U36b4* measured by quantitative real-time PCR. Data represent mean ± SEM for 3 males and 3 females per time point and light condition. Rhythmicity was determined by JTK_Cycle analyses; *P^JTKCycle^ <0.05, **P^JTKCycle^ <0.01, ****P^JTKCycle^ <0.0001. (**C**,**E**) Proteins detected by immunoblot. Each lane on the Western blot represents a sample prepared from a unique animal. Representative images were taken from n = 6 biological replicates.

The expression of clock-controlled genes was also impacted by CJL. *Wee1* is a critical regulator of the G2/M transition of the cell cycle that is under circadian control via BMAL1-CLOCK activation that targets E-box elements in the *Wee1* gene promoter (*29*). *Wee1* expression was strongly influenced by the time of day in healthy lungs and was robustly impacted by CJL (Fig.1D). CJL also profoundly altered the diurnal expression of *Cxcl5* (Fig.1D), a key chemokine for recruiting neutrophils to the lung upon bacterial and viral exposures (*30-32*) that is regulated by REV-ERBα and REV-ERBβ (*33*). However, expression of the cell cycle regulators *p53* and *p21*, and oncogenic transcription factor *c-Myc*, each of which can be regulated by circadian clock factors (*34-36*), were not affected by CJL in lung tissue (Fig.1D). Although p53 and c-MYC are regulated post-translationally by clock components (*35, 36*) and exhibit rhythmic expression that is impacted by exposure to altered lighting conditions in mouse thymus (*23*), we did not observe any clear effect of CJL on the levels of these proteins in the lungs of healthy mice (Fig. 1E).

In an earlier study of mice with mammary tumors in the FVB genetic background, chronic disruption of light exposures dramatically increased weight gain (*37*). In contrast, we found that CJL had no impact on body weight in healthy C57BL/6J mice (fig. S2A). Interestingly, we observed CJL-induced differences in rhythmic corticosterone levels in serum of female mice only, suggesting a more rapid light entrainment for corticosterone secretion in males and sexual dimorphism in hormonal responses (fig. S2B). In a separate group of mice that were housed first in 12:12LD for 2 weeks and then in CJL for 13 weeks with access to a running wheel, we confirmed disruption of rhythmic behavior upon CJL; while the locomotor activity was mostly consolidated within the dark phases, the pattern and amplitude of activity during the dark hours dramatically changed after only one week of CJL (fig. S3A-E).

### Experimental Chronic Jetlag Increases Kras^G12D^-driven Lung Tumor Burden

In order to investigate molecular mechanisms related to light-induced circadian disruption in lung tumorigenesis, we used a genetically-engineered mouse model of Non-Small Cell Lung Cancer in which tumor formation is initiated by expression of oncogenic *Kras*^*G12D*^ in a small number of lung cells (*Kras*^*LSL-G12D/+*^ mice, also known as K mice) – a model established to recapitulate many of the clinical features of naturally occurring KRAS-driven lung cancer (*38*). Male and female K mice were first infected intratracheally with lentivirus expressing CRE recombinase under the control of the *Ubc* promoter to induce tumorigenesis, and five weeks later were placed in 12:12LD or CJL conditions, ensuring that tumor initiation rates and the first 5 weeks of tumor growth were under standard conditions before the onset of CJL, as previously reported (*39*). At 25 weeks post-infection (20 weeks in CJL), we observed a striking 60% increase in tumor burden in K mice housed in CJL conditions compared to those that had remained in 12:12LD (Fig. 2A,B). This was attributed to an increase in the number (Fig. 2C) and not the size (Fig. 2D) of tumors, suggesting that CJL impacts early events in tumor progression in this model. Moreover, there was no difference in the spectrum of tumor grades assessed by histopathology (*38*) between the two groups, with most of the tumors being grade 2 adenomas (Fig. 2E). There was no impact of CJL on overall survival (Fig. 2F), indicating that additional factors precipitating death in K mice appeared stochastically in both light conditions. An earlier study demonstrated that CJL increased tumor burden in *K-ras*^*LSL-G12D/+*^;*p53*^*flox/flox*^ (KP) mice, but to a much lesser extent than we observed in K mice (*39*). Consistent with this, we did not observe any effects of CJL on lung tumor burden, numbers, grading, or overall survival in KP mice (fig. S4), which harbor simultaneous activation of oncogenic K-RAS and deletion of tumor suppressor protein p53, and thus experience more rapid and severe tumor progression. Technical differences that resulted in overall lower tumor burden in the KP mice studied by Papagiannakopoulos et al., may have enabled them to observe an effect of CJL where we did not.

**Fig. 2.**
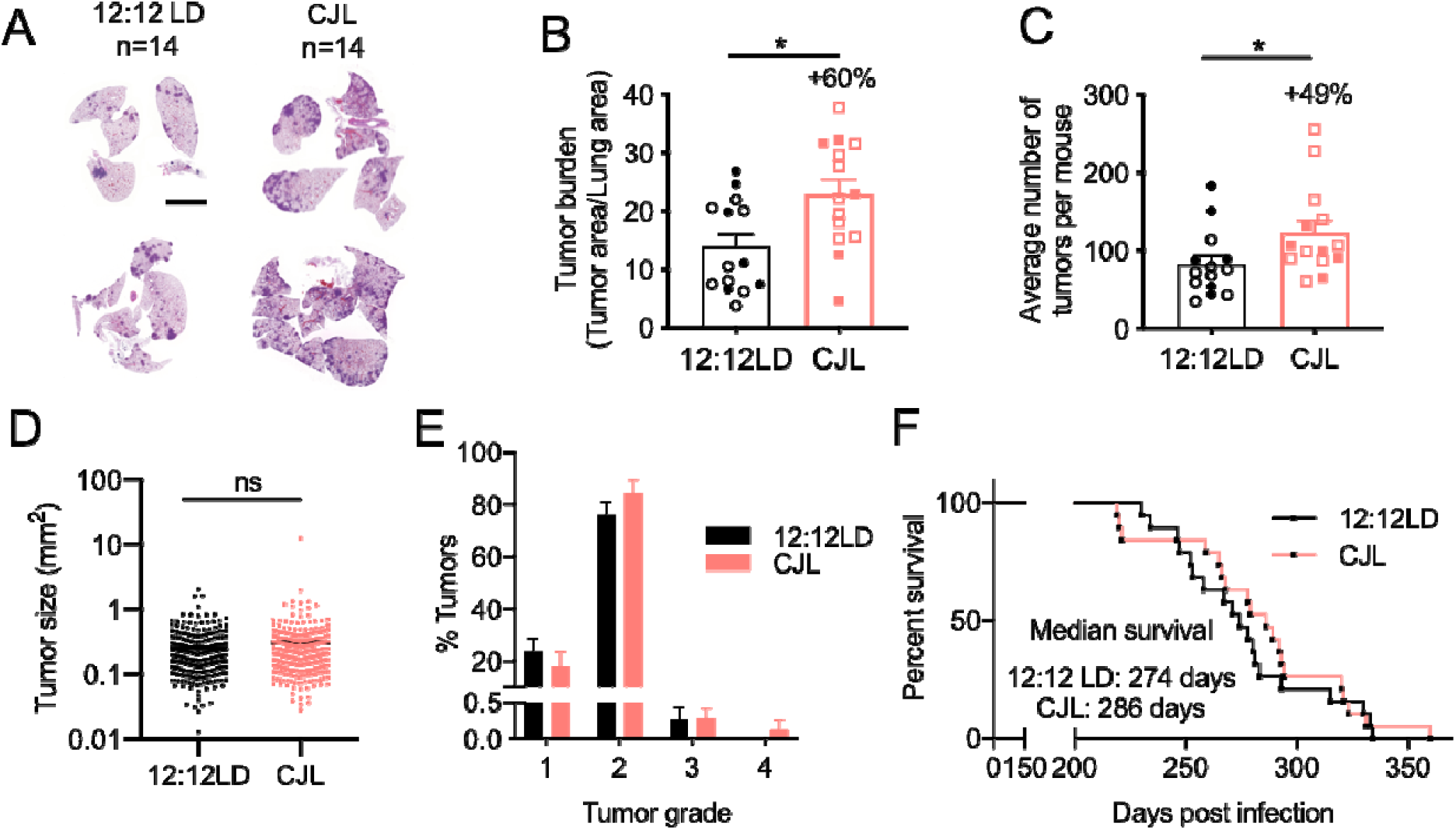
Chronic jetlag (CJL) causes an increase in tumor burden in K mice but has no impact on survival. Five weeks post-infection with lentivirus-Cre, K mice were placed in either 12:12LD or CJL for 20 weeks (**A-E**) or until signs of distress (**F**). (**A**) Representative H&E-stained sections at endpoint; scale bar, 5000 µm. Tumor burden (**B**), numbers (**C**), size (**D**) and grade (**E**) were assessed from H&E sections. Column data represent mean ± SEM. Values for individual animals (**B**,**C**,**E**) or tumors (**D**) are plotted. **(B**,**C**) Clear and filled symbols represent males and females respectively. *P <0.05 by Mann-Whitney test. **(F)** Kaplan-Meier survival analysis for K mice placed in 12:12LD (n = 19) or CJL (n = 19) conditions.

### c-MYC levels in K mice do not explain increased tumor burden upon CJL

The c-MYC oncoprotein has been shown to play a crucial role in the growth of KRAS-driven lung tumors (*40*) and c-MYC accumulation in mouse thymus exhibits a daily rhythm and is robustly elevated at all times of day after exposure to a single shift of light exposure (*23*). Furthermore, genetic deletion of *Bmal1* or loss-of-function of PER2 enhanced c-MYC accumulation in *Kras*^*G12D*^-driven lung tumors (*3*). The circadian transcriptional repressor CRY2 can recruit phosphorylated substrates, including c-MYC (*36*), to the SCF^FBXL3^ ubiquitin ligase, thereby promoting their ubiquitination and proteasomal degradation. We thus expected that CJL could promote tumorigenesis by perturbing the expression of CRY2, resulting in aberrant accumulation of c-MYC. Unexpectedly, CJL resulted in significantly reduced accumulation of c-MYC in *Kras*^*G12D*^-driven lung tumors as assessed by immunohistochemistry and Western blotting (fig. S5). Thus, while aberrant stabilization of c-MYC may contribute to enhanced cell growth and transformation caused by deletion or suppression of circadian clock components (*3, 36*), it does not appear to play a major role in the enhanced tumorigenesis caused by circadian disruption of environmental light exposure in the context of KRAS-driven lung cancer.

### CJL further disrupts an already dysregulated clock machinery in tumors from K mice

To identify mechanisms potentially underlying the pro-tumorigenic effect of CJL in K mice, we compared the transcriptional programs in tumors and total lung from K mice housed in normal or CJL conditions. Up to 12 individual tumors were collected from each animal at either ZT9 or ZT21 and the remaining lung tissue was also collected for subsequent analyses. We chose these two time points as they represent the trough and peak, respectively, of expression of the core clock component, *Bmal1*, in healthy lung under normal light conditions. We sequenced RNA prepared from 3 tumors per animal and two mice per time point and lighting condition (Fig. 3A). To gain unbiased insight into transcriptional networks perturbed by CJL, we used differential expression analysis (DESeq2) (*41*). The ‘whole lung’ samples also contain several tumors, but the tumor tissue represents a smaller fraction of the whole compared to the tumor samples. Pathway enrichment analysis confirmed up-regulation of KRAS signaling in resected tumors compared to whole lung samples for all conditions combined (fig. S6A). When looking at each time point individually for tumors, we identified 53 and 85 genes differentially expressed between 12:12LD and CJL at ZT9 and ZT21, respectively, with 20 genes differentially expressed at both time points (Fig. 3B,C, fig. S6B). The expression changes for these 20 genes were inverted between ZT9 and ZT21 (fig. S6B), suggesting that their expression is under circadian control. Core clock genes and highly rhythmic transcription factors *Tef* and *Dbp* were among the most differentially expressed genes in all groups, and not surprisingly DAVID analyses highlighted biological circadian rhythms and rhythmic processes in the cluster with the highest enrichment score for tumor groups at each time point (fig. S6C). Transcript analyses by qPCR of different tumors and additional lung samples validated these findings, demonstrating significant variations in *Bmal1, Per2* and *Rev-erbβ* mRNA between tumors from K mice housed in 12:12LD and CJL conditions (Fig. 3D). Notably, rhythmic expression of core clock genes was retained in tumors from mice housed in the 12:12LD standard light condition, but the amplitude of some clock gene expression such as *Per2, Cry2* and *Rev-erbα* was reduced in tumor samples compared to whole lungs (Fig. 3D). This is consistent with several studies demonstrating dampening of circadian rhythms in tumors, mediated by oncogenic factors such as c-MYC or RAS (*42, 43*). Similar results were observed at the protein level, with a particularly dramatic reduction in REV-ERBα at the peak of its normal expression (ZT9) in tumors compared to total lung from K mice housed in 12:12LD (Fig. 3E). REV-ERBα was not detected at the trough of its expression (ZT21) in either tumors or total lung, indicating that rhythmicity of REV-ERBα was probably retained in tumors but with a greatly diminished amplitude. Moreover, REV-ERBα protein levels were very low under CJL conditions in both tumors and total lung samples at these two time points, but given its very high amplitude of expression, we cannot exclude the possibility that CJL caused a shift in the phase of REV-ERBα.

**Fig. 3.**
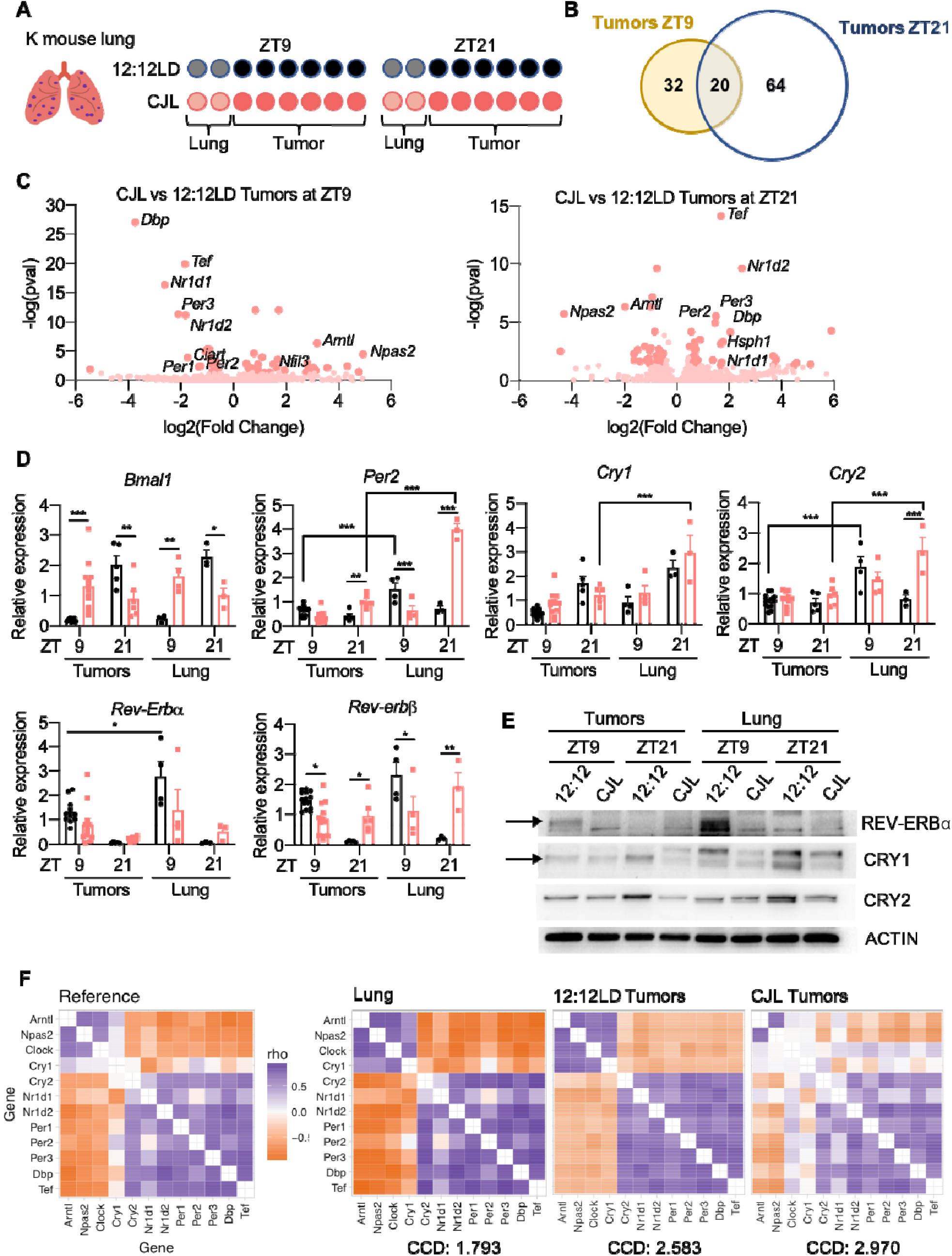
Chronic jetlag (CJL) further disrupts an already dysregulated clockwork in tumors from K mice. Five weeks post-infection with lentivirus-Cre, K mice were placed in either 12:12LD or CJL for 20 weeks. (**A**) For RNA-sequencing, 3 tumors per animal and two mice per time point and light conditions were used. (**B**) Plots indicating the numbers of differentially expressed genes between 12:12LD and CJL by DESeq2 analyses for each condition, with adj. p-value<0.05 and fold change >+/- 1.4 cut-offs. (**C**) Volcano plots of differentially expressed genes between 12:12LD and CJL by DESeq2 analyses for tumors collected at ZT9 and ZT21. (**D**) Gene expression normalized to *U36b4* measured by quantitative real-time PCR. Data represent mean ± SEM; n = 14 and 12 for tumors collected at ZT9 in 12:12LD and CJL respectively, n= 5 and 6 for tumors collected at ZT21 in 12:12LD and CJL respectively and n=3-4 for lung samples. *P <0.05, **P <0.01 and ***P <0.001 by two-Way ANOVA, post hoc Bonferroni test. (**E**) Proteins detected by immunoblot. Tumors and lungs for each light condition and time point on the blot were from the same animal. Representative images were taken from n = 3 biological replicates. (**F**) Heatmaps of Spearman correlation between each pair of the 12 clock genes and corresponding clock correlation distance (CCD; relative to the mouse reference) in murine healthy lung from previously published data set GSE54651, or in tumors from 12:12LD or CJL conditions.

To further assess whether CJL disrupts the clockwork in tumors from K mice, we used the clock correlation distance (CCD) algorithm, which infers the regularity of circadian clock progression in a group of samples based on the correlated co-expression of 12 clock genes (*44*). A higher CCD score indicates a more profound disruption of circadian rhythmicity. Originally, the CCD method was designed to evaluate circadian clocks in human cancer and revealed that clock gene co-expression in tumors is consistently perturbed compared to matched healthy tissue. Because our whole lung samples contain both healthy lung tissue and tumors, we compared the CCD score from our 12:12LD tumors (*n* = 12) - calculated using our data acquired by sequencing RNA - to that calculated using published data from murine healthy lung (*45*). This analysis revealed that the CCD score for healthy mouse lung was lower than the CCD score for tumors from K mice housed in 12:12LD. Moreover, we found that the CCD score for tumors from CJL-housed mice was higher than the CCD score for tumors collected from control animals (Fig. 3F), although the difference between groups is not statistically significant, likely due to limited numbers of samples. Together, these results strongly suggest a dysregulation of circadian clock progression in tumors from K mice compared to healthy lung tissue under normal light condition, and that exposure to CJL further perturbs an already disrupted clock within KRAS^G12D^-driven lung tumors.

### HSF1 signaling is upregulated in response to CJL in K mice

Because we were primarily interested in identifying gene networks that were consistently impacted by circadian disruption within both tumors and lungs without time of sample collection as confounding factor, we searched for transcripts that were differentially expressed between all the samples (tumors + lungs) collected from mice housed in control 12:12LD lighting conditions compared to those housed in CJL lighting conditions. This comparison revealed that genes encoding various heat shock proteins (HSPs) are upregulated in samples from CJL-exposed K mice (Fig. 4A). The same analysis including only tumor samples gave very similar results (fig. S7A). The expression of HSPs is primarily activated by the heat shock response-associated transcription factor heat shock factor 1 (HSF1) in response to pathologic insults that disrupt cytosolic proteostasis, including modest changes in temperature and oxidative stress (*46, 47*).

**Fig. 4.**
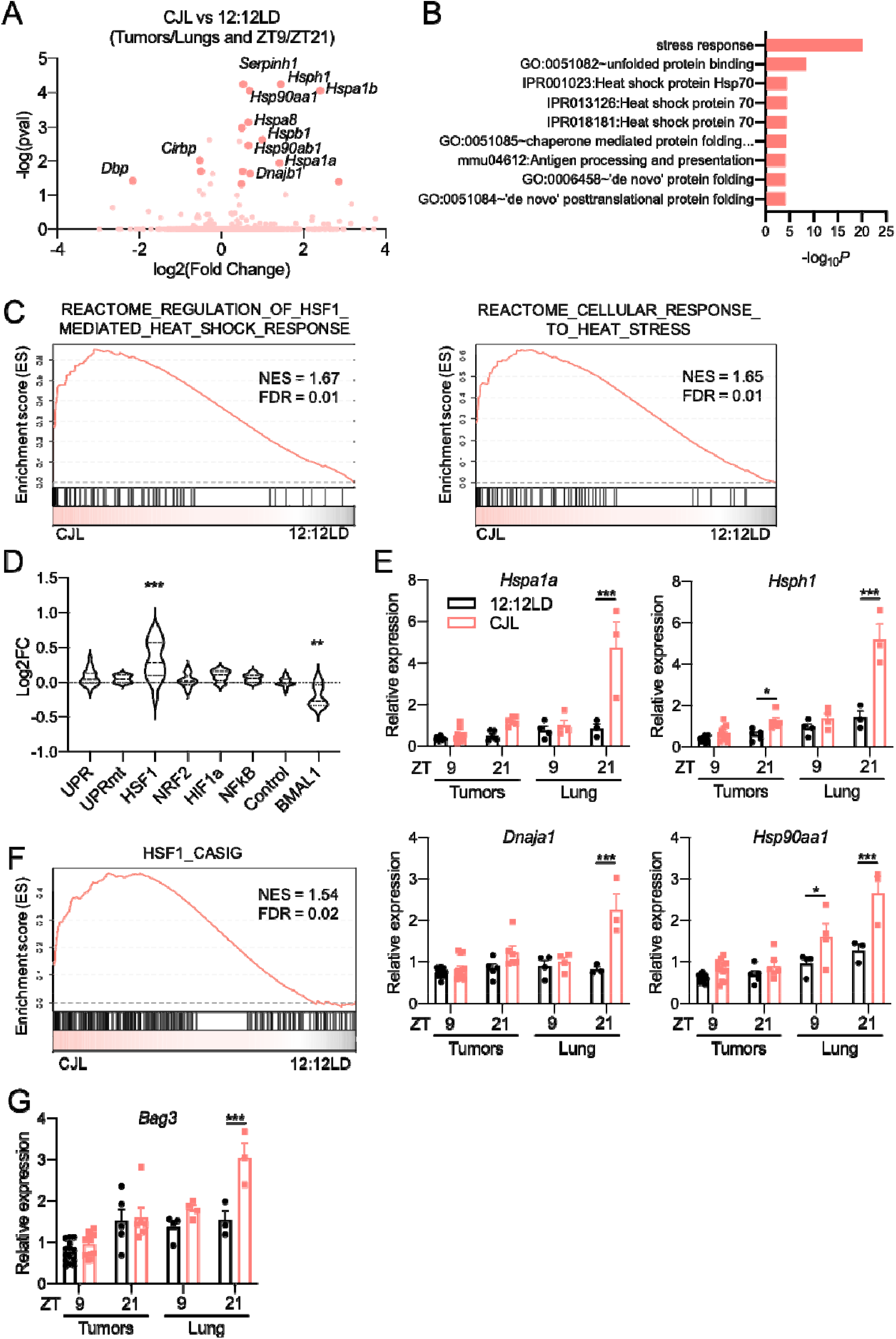
CJL enhances expression of the HSF1-mediated heat shock response and cancer signature in Kras^G12D^-driven lung tumor model. Five weeks post-infection with lentivirus-Cre, K mice were placed in either 12:12LD or CJL for 20 weeks. **(A)** Volcano plots of differentially expressed genes between 12:12LD and CJL by DESeq2 analyses for all samples (tumors+lungs), taking time of collection (ZT9/21) as confounding factor. **(B)** DAVID analyses on the differentially expressed genes by DESeq2 in all samples between 12:12LD and CJL, taking time of collection as confounding factor. Only terms with FDR< 0.25 are shown. **(C)** GSEA plots for the cellular response to heat stress and HSF1-mediated heat shock response reactome gene sets applied to samples (lungs+ tumors) from CJL vs 12:12LD housed K mice. **(D)** Activation of stress response pathways by CJL in lungs and tumors, independently of collection time, revealed by grouped fold change for transcripts established as selective targets of each stress pathways (Grandjean et al., 2019) or “BMAL1-pathway” (Shilts et al., 2018). **P < 0.01, **** P < 0.0001 by one-Way ANOVA with Dunnett’s multiple comparison test. **(E**,**G)** Gene expression normalized to *U36b4* measured by quantitative real-time PCR; T = tumors, L = lungs. Data represent mean ± SEM; n = 14 and 12 for tumors collected at ZT9 in 12:12LD and CJL respectively, n= 5 and 6 for tumors collected at ZT21 in 12:12LD and CJL respectively and n=3-4 for lung samples. *P <0.05, **P <0.01 and ***P <0.001 by two-Way ANOVA, post hoc Bonferroni test. **(F)** GSEA plot for the gene set representing the HSF1-Cancer signature network applied to samples (lungs+ tumors) from CJL vs 12:12LD housed K mice.

HSPs function to enhance proteostasis capacity of the cell and prevent the pathologic accumulation of potentially toxic protein aggregates (*48*). Accordingly, DAVID analysis pointed to stress response and response to unfolded protein in the cluster with the highest enrichment score (Fig. 4B, fig. S7B). Strikingly, all of the transcripts that are significantly elevated in samples from mice exposed to CJL are known transcriptional targets of HSF1 (*49*). Gene Set Enrichment Analysis (GSEA) provided further support for the idea that CJL leads to an elevated HSF1-mediated heat shock response upon CJL in these samples (Fig. 4C). Moreover, this CJL-driven activation of HSF1 in K mice appeared to be selective, as we did not observe increased expression of genes regulated by other stress-responsive signaling pathways (Fig. 4D) (*50*). Consistent with dysregulation of clock gene expression measured by qPCR (Fig. 3D), we also observed a significant decrease in expression of a set of previously defined BMAL1 target genes (*44*) (Fig. 4D, right). Transcript analyses by qPCR from different tumors and additional lung samples validated these findings and highlighted a more pronounced upregulation of HSF1 target gene expression at ZT21 in whole lungs from mice exposed to CJL compared to normal light conditions (Fig. 4E).

HSF1 broadly influences tumor biology (*21*). Notably, HSF1 activates a distinct transcriptional program in malignant cells, dubbed the HSF1 cancer signature, or HSF1-CaSig (*51*). We created a gene matrix based on the HSF1-CaSig defined in (*51*) and used GSEA to show that samples from CJL-exposed mice also exhibit robustly enriched expression of the HSF1-CaSig network compared to lungs and tumors from control mice (Fig. 4F). BCL2-associated athanogene 3, *Bag3*, is a molecular chaperone and HSF1 target gene that is part of the HSF1-CaSig network, and is involved in apoptosis evasion (*52*). qPCR revealed that *Bag3* is significantly upregulated in whole lung collected at ZT21 from K mice exposed to CJL compared to normal light conditions (Fig. 4G), mirroring changes measured for other HSF1 target genes, like *Hspa1a* (Fig. 4E).

Interestingly, applying GSEA to a previously published gene expression profiling dataset of Kras^G12V^-driven lung hyperplasia and normal murine lung cells (*53*) showed that both the HSF1-mediated heat shock response and HSF1-CaSig gene sets were significantly enriched in hyperplastic lesions compared to normal lung cells (fig. S7C). However, these gene sets were not significantly enriched in frank adenocarcinoma compared to hyperplastic lesions in the same dataset (fig. S7D). This suggests that activation of HSF1 signaling occurs at early stages of KRAS-driven lung cancer development.

### Rhythmic HSF1 nuclear accumulation and transcriptional activity is perturbed by CJL

Previous studies show that nuclear HSF1 levels fluctuate daily in the liver of mice and chipmunks in phase with body temperature rhythms, and that HSF1 acts as a circadian transcription factor (*54, 55*). To determine if this was also the case in lung tissue and to assess the impact of CJL exposure on this regulation, we measured HSF1 protein levels in lung nuclear extracts over 24 hours from C57BL/6J mice housed in 12:12LD or CJL conditions (Fig. 5A,B). Lung nuclear HSF1 protein levels exhibited robust diurnal oscillations, peaking during the dark phase, under normal light conditions. Interestingly, upon CJL exposure, the amplitude of this rhythm was dampened but the phase was retained (or only slightly shifted) leading to enhanced accumulation of nuclear HSF1 at the beginning of the light phase. In contrast, total HSF1 protein levels exhibited a moderate rhythm with lower amplitude and different phase than nuclear HSF1 levels (fig. S8). The expression profiles of HSF1 target genes from a different cohort of mice mirrored diurnal HSF1 nuclear localization (Fig. 5C), with enhanced expression upon CJL from the end of dark phase through the beginning of the light phase. These results were also consistent with increased expression of HSF1 target genes that we observed at ZT21 in K mice exposed to CJL (Fig. 4E). These findings indicate that CJL perturbs homeostatic regulation of HSF1 transcriptional activity in the lung, which could lead to enhanced tumor initiation in combination with other oncogenic factors.

**Fig. 5.**
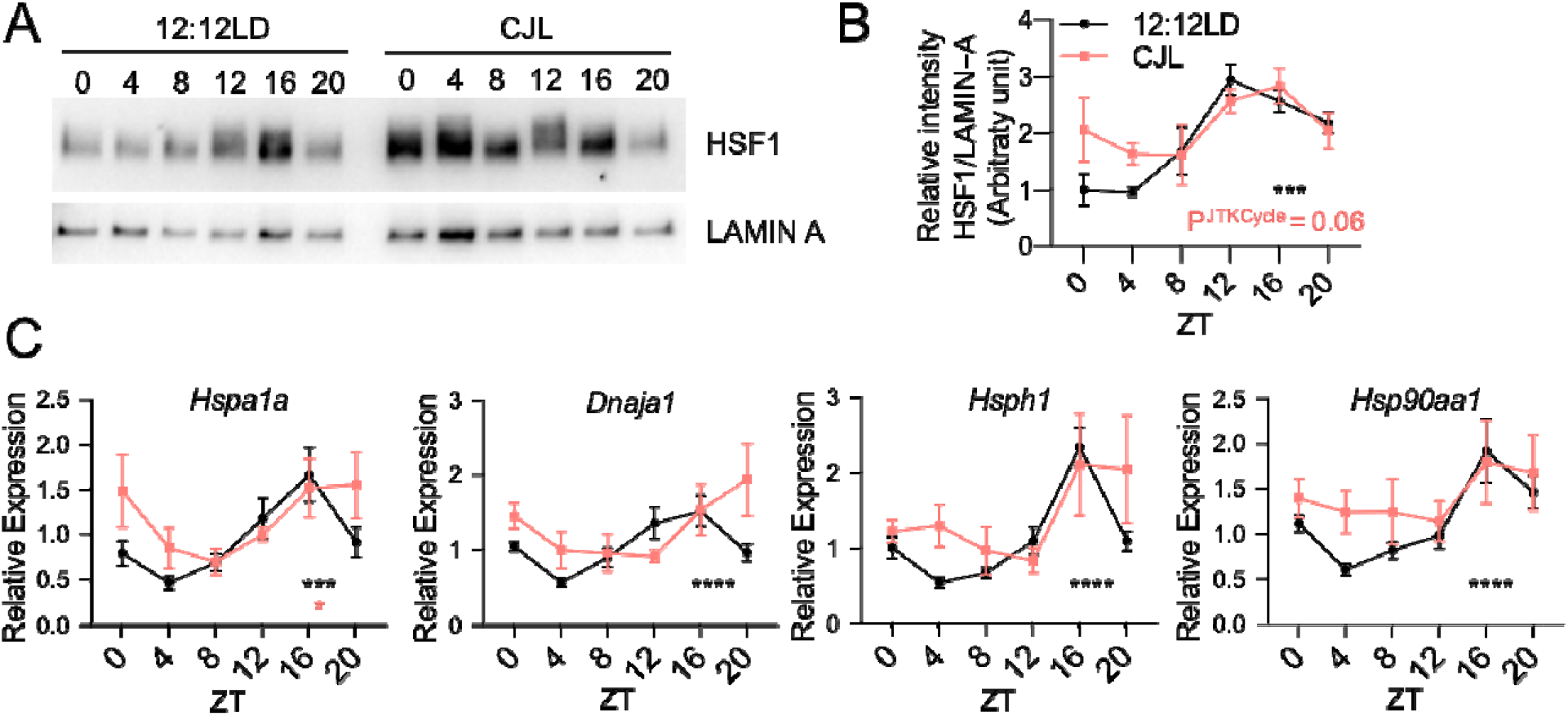
Time-regulated HSF1 nuclear accumulation and transcriptional activity are perturbed upon CJL exposure. C57BL/6J mice were housed in 12:12LD or CJL for 8-12 weeks. Lung tissues were collected at the indicated times (hours after lights on) on Day 1 of the schedule shown in Fig. 1A. **(A)** Proteins from lung nuclear extracts detected by immunoblot. Each lane on the Western blot represents a sample prepared from a unique animal. Representative images were taken from *n* = 3 biological replicates. **(B)** Quantitation of (A). Rhythmicity was determined by JTK_Cycle analyses; ***P^JTKCycle^ <0.001. **(C)** Gene expression normalized to *U36b4* measured by quantitative real-time PCR. Data represent mean ± SEM for 3 males and 3 females per time point and light condition. Rhythmicity was determined by JTK_Cycle analyses; *P^JTKCycle^ <0.05, ***P^JTKCycle^ <0.001, ****P^JTKCycle^ <0.0001.

### HSF1 signaling affects human KRAS-mutant lung cancer

To gain insight into the potential for HSF1 to influence human lung adenocarcinoma, we treated human lung cancer cell lines with a novel Direct Targeted HSF1 InhiBitor (DTHIB) that has been shown to stimulate degradation of nuclear HSF1 and suppress the growth of prostate cancer xenografts (*56*). We confirmed the potency of DTHIB for suppressing HSF1 transcriptional activity in HEK293T cells expressing a Heat Shock Element (HSE)-Luciferase reporter in which HSF1 is stimulated with the activating ligand A3 (*57*) (fig. S9A). We found that inhibiting HSF1 with DTHIB slowed the growth of two human lung adenocarcinoma (LUAD) cell lines harboring heterozygous KRAS^G12D^ mutations (A-427 and SK-LU-1 cells) in a dose-dependent manner (Fig. 6A,B; fig. S9B,C for later treatment). We confirmed significant reduction in the protein levels of HSF1 and downstream chaperone DNAJB1 (HSP40) in A-427 cells treated with 5 µM DTHIB for 48 hours, confirming compound activity in this model (Fig. 6C,D). We did not detect a significant change in KRAS^G12D^ protein level upon DTHIB treatment. However, two downstream effectors of RAS signaling (phosphorylation of ERK1/2 and accumulation of c-MYC) were significantly decreased (Fig. 6C,D), suggesting impairment of RAS signaling in these cells upon pharmacologic inhibition of HSF1. Altogether, these results indicate that HSF1 is an important contributor to cellular proliferation in two cell models of KRAS-driven LUAD.

**Fig. 6.**
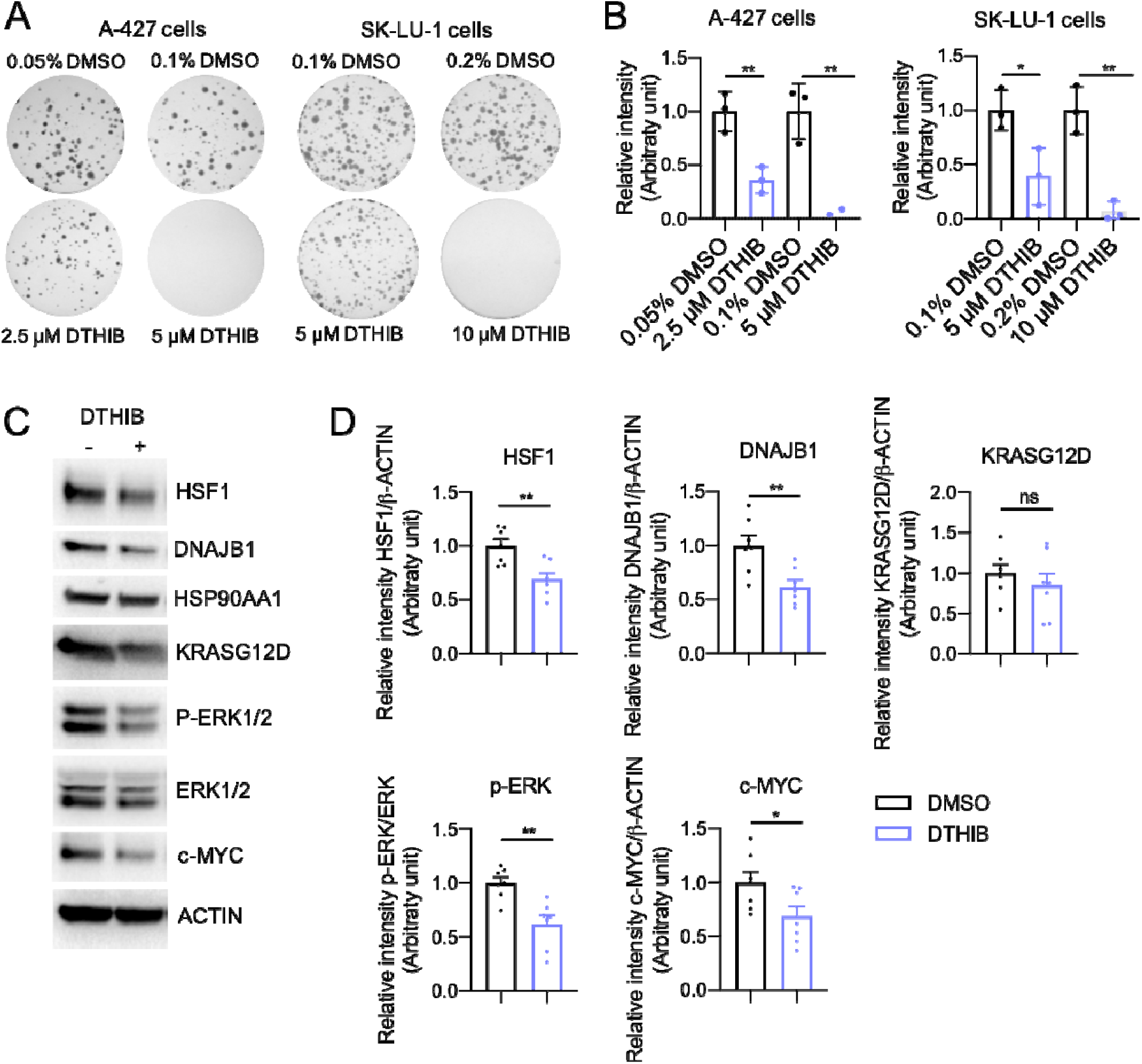
HSF1 signaling influences human KRAS-mutant LUAD cell growth. **(A)** Representative images of crystal violet stained colonies formed by A-427 or SK-LU-1 cells treated with DTHIB or vehicle DMSO 2 days after seeding, for 14 days. **(B)** Quantification of (A) from three biological replicates. Each condition was compared to controls that were plated in wells on the same plates. Bars represent mean ± SD, **P < 0.01 by student t-test. **(C)** Immunoblot of A-427 cells treated with 5 µM DTHIB or 0.1% DMSO for 48h. Representative of *n*=7. **(D)** Quantitation of (C).

## Discussion

Several molecular hypotheses have been proposed to explain the increased cancer risk associated with circadian disruption. Studying the impact of circadian disruption in genetically engineered mouse models of cancer enables us to identify molecular changes that occur in response to chronic jetlag and investigate their contributions to enhanced cancer risk in a controlled environment. Here, we show that circadian disruption promotes lung tumorigenesis in K mice, a genetically-engineered mouse model in which tumor formation is initiated by expression of oncogenic *Kras*^G12D^ in a small number of lung cells that recapitulates many of the clinical features of naturally occurring KRAS-driven lung cancer (*38, 58*). This provides additional evidence of the impact of circadian disruption on lung cancer as previously reported in a related but more severe lung cancer mouse model (*3*). Interestingly, the CJL protocol used affects the number of tumors detected but not their size. This indicates that circadian disruption impacts early events in Kras^G12D^-driven tumor progression or prevents regression of initiated tumors rather than enhancing the growth of well-established tumors. We further demonstrated that HSF1 signaling is significantly elevated in mice exposed to altered light/dark cycles, revealing a novel mechanism of action that likely contributes to increased tumor formation in response to circadian disruption.

The CJL protocol used in our studies, consisting of repeated light advances and mimicking the effects of rotating shift work or frequent eastbound transmeridian flights, has been previously shown to cause severe perturbations in rest-activity cycles and body temperature (*3, 25, 26, 59*). By housing mice in constant darkness for two days after ten days of this CJL protocol exposure (to avoid any masking effect of light on circadian rhythmic patterns) previous work has demonstrated that this protocol caused dysregulation of circadian rhythms of gene expression in the SCN and peripheral organs (*25, 26, 60*). In our studies, mice were kept in their respective light schedule and tissues were sampled when all mice experienced the same light exposures for at least 24 hours prior to sample collection. In that way, we aimed to assess the effects of CJL in a normal light exposure context, which is more representative of what humans experience after shift work or frequent eastbound flights. As expected, we found that peripheral clocks cannot adjust their timing rapidly enough to maintain synchrony with the shifting of the environment. Interestingly, it appeared that some clock genes, including *Per2*, remained rhythmic after CJL exposure while others (e.g. *Bmal1*) did not, consistent with prior work indicating that *Per2* is more sensitive to entrainment signals in peripheral tissues than other core clock genes (*61*).

Our analysis of gene expression in tumors and tumor-bearing lung tissues from animals housed in standard or CJL conditions demonstrated that the clock machinery was highly disrupted by CJL in both tumors and lung tissue. Accumulating evidence reveals that circadian clock components play critical roles in several hallmarks of cancer, including cell proliferation, DNA damage and repair, and cell death (*15*), suggesting that disruption of cellular circadian rhythms within tumors could contribute to the detrimental impact of irregular light exposure in cancer. In support of this idea, previous work established that K mice, the same *Kras*^G12D^-driven NSCLC mouse model that we used, develop a greater tumor burden when the core circadian clock component *Bmal1* is deleted specifically within tumors (*3*). Conversely, previous studies have shown that circadian functions in cancer cells are compromised or deregulated (*44*), in some cases due to high expression of oncogenic c-MYC (*42*) or RAS (*43*). However, circadian disruption does not seem to be a universal feature of cancer, because some cancer cells, such as melanoma, Acute Myeloid Leukemia cells and patient-derived cancer stem cells (CSCs) of glioblastoma harbor an intact circadian clock despite their highly tumorigenic and metastatic potential (*62-64*).

Here, we demonstrate that chronic circadian disruption *in vivo* enhances the expression of HSF1 target genes. HSF1 promotes expression of heat shock proteins (HSPs) to protect the proteome, allowing cells to survive diverse proteotoxic stresses. Over the past decade, it has become clear that HSF1 activity is exploited by cancer cells to overcome diverse stresses and intrinsic and extrinsic demands (*65*). High levels of HSF1 and HSF1-regulated HSPs have been measured in different types of cancers and are negatively correlated with prognosis in patients (*66, 67*), including those with NSCLC (*68*). Accordingly, in various human cancer cell lines and murine cancer models, deletion of HSF1 markedly reduces growth, survival and metastatic potential (*21, 51, 69-72*), whereas its overexpression enhances the malignant phenotypes of xenografted human melanoma cells *in vivo* (*73*). Several mechanisms have been proposed to contribute to the role of HSF1 in supporting malignancy, including regulation of HSPs expression and regulation of an unique transcriptional program activated by HSF1 in cancer cells dubbed the HSF1 cancer signature (HSF1-CaSig) (*51*). Notably, the HSF1-CaSig was up-regulated in tumor-bearing K mouse lung upon CJL exposure (Fig. 5F). Regulation of cancer cell proteostasis by HSPs is an important feature for cancer cell survival and proliferation (*74*) and recently, HSPs have drawn increased attention as potential targets in cancer, especially given the role of such stress proteins in promoting resistance to conventional therapies (*75*). Specifically, growing evidence of correlation between *Hsps* expression profile and degree of differentiation and staging of lung tumors suggest that these proteins could be considered as therapeutic targets and biomarkers for lung cancer patient management (*76, 77*). Indeed, mutant oncoproteins, such as KRAS^G12D^, may depend on HSF1-dependent regulation of HSPs to enable folding and to maintain full activity. Although we did not detect a significant change in KRAS^G12D^ protein level upon pharmacological inhibition of HSF1, its activity appeared to be impaired as measured by significant down-regulation of the phosphorylation levels of its downstream effectors ERK1/2. These findings are consistent with previous observations, in other contexts, that HSF1 supports transformation and tumorigenesis via activation of oncogenic RAS signaling (*71, 78*).

Our work also adds to a growing body of evidence indicating robust circadian regulation of HSF1 activity, both in healthy and tumor-bearing lung tissue. As in the liver (*54*), HSF1 nuclear accumulation in the lung is dependent on the time of day, peaking at night under normal light conditions. Daily rhythmic nuclear accumulation of HSF1 is associated with rhythmic expression of *Hsps*, which peak in the middle of the dark phase, around ZT16. Exposure to CJL disrupted these rhythms mainly by preventing the reduction of their expression that is seen between ZT20 and ZT4 under normal conditions. Interestingly, fluctuations in body temperature have been shown to act as a major entrainment factor for rhythmic HSF1 activity (*54*), and compared to day-shift nurses, night-shift nurses exhibited significant differences in peripheral skin temperature, with notably higher minimum temperature but unchanged maximum temperature (*79, 80*). Therefore, activation of HSF1 due to abnormal changes in body temperature could be a key component in the connection between shift work and cancer risk.

Numerous cell-autonomous and systemic mechanisms are susceptible to alteration upon circadian disruption and can influence tumorigenesis. In this work, we revealed that circadian disruption impacts early events in tumor formation in a KRAS-driven mouse model of lung adenocarcinoma. Further, we demonstrated that HSF1 signaling in the lung and lung tumors is dysregulated by exposure to altered environmental lighting schedules. HSF1 has been shown to support tumorigenesis in myriad ways (*81*), suggesting that the enhanced HSF1 activity that we observed in response to circadian disruption could play an important role in increased tumor formation. Additional investigation is needed to determine whether the chronic elevation of HSF1 signaling that we measured in lungs in response to circadian disruption occurs early in the disease process, whether it is present in other anatomical locations, and whether HSF1 is required for increased tumorigenesis in response to circadian disruption. We demonstrated that a novel direct HSF1 inhibitor (DTHIB), previously shown to potently attenuate tumor progression in therapy-resistant prostate cancer models (*56*), reduced growth of two different human KRAS-driven lung cancer cell lines. To further define the determinants of susceptibility to growth inhibition by DTHIB, it will be necessary to examine additional cell lines and to investigate the impact of genetic manipulation of HSF1, KRAS, and other factors on growth inhibition by DTHIB. Of particular relevance for connecting HSF1-related therapeutic opportunities to circadian disruption, an HSP90 inhibitor reduced the growth of mouse melanoma in a time-of-day–specific manner, which depended on an intact core clock system in the tumors (*82*). Our findings described herein demonstrate that HSF1 and its downstream effectors, the HSPs, are potential therapeutic targets for mitigating cancer risk among populations exposed to chronic circadian disruption, such as shift workers.

## Materials and Methods

### Mouse Models

C57BL/6J mice were purchased from the Scripps Research breeding colony at six weeks of age. They were group housed except when given voluntary access to running wheels, in which case they were singly housed in running wheel cages. Genetically engineered mouse models, Kras^LSL-G12D/+^ (K) and Kras^LSL-G12D/+^;p53^flox/flox^ (KP), all in pure C57BL/6J background, were obtained from the Jackson Laboratory, and have been previously described (*38, 58*). When they were between eight and ten weeks old, mice were infected intratracheally with lentivirus containing Cre recombinase, lenti-Cre (PGK-Cre, gift from Tyler Jacks), at a viral titer of 5 × 10^5^ plaque forming units (PFU) per mouse according to the previously established protocol (*38*). All experiments utilized both female and male mice. They were given ad libitum access to normal mouse chow and water. Sacrifices during the dark phase were carried out under red light. All animal care and treatments were in accordance with Scripps Research guidelines for the care and use of animals, and approved by the Scripps Research Institutional Animal Care and Use Committee (IACUC) under protocol #10-0019.

### Chronic Jetlag (CJL) Conditions

Mice were randomly placed into standard light conditions (12:12LD) or chronic jetlag (CJL) consisting of an 8-hour light phase advance repeated every 2 or 3 days (*3, 23, 60*). For GEMM studies, mice were housed in these light conditions 5 weeks post infection with lenti-Cre, for 10 or 20 weeks for KP and K mice respectively, or until showing signs of distress for the survival studies.

### Running Wheel Activity Analysis

C57BL/6J mice were single housed and given access to running wheels with ad libitum access to food and water for seceral weeks, under specific light conditions as indicated in the figure legends. Voluntary running wheel activity was analyzed with ClockLab (Actimetrics) using digital recordings of wheel rotations.

### Histology and tumor burden analyses

Mice were euthanized by carbon dioxide asphyxiation. Lungs were inflated through the trachea with 4% paraformaldehyde (PFA), fixed overnight, transferred to 70% ethanol and sent to the Rodent Histopathology Core facility at Harvard Medical School for subsequent paraffin-embedding and sectioning at a thickness of five micrometers. Sections were stained with haemotoxylin and eosin (H&E) for pathological examination. Histopathological grading of tumors and quantification of tumor numbers were performed with the assistance of Dr. Roderick Bronson, histopathologist at Harvard Medical School. Tumor size of each individual tumor was measured from H&E stained sections using morphometric analysis in Panoramic viewer software (Perkin Elmer). Tumor burden, calculated as a percentage of tumor area per total lung area per mouse, was quantified from H&E stained sections using a Nuance automated spectral imaging system (Inform v2.1 software, Cambridge Research and Instrumentation). In brief, the Trainable Tissue Segmentation method was trained to identify tumor, normal lung, vessel and space. This program was then applied to all H&E images, and each of the resulting mapped images was then screened to verify that accurate tissue segmentation had occurred.

### Immunohistochemistry

Slides were deparaffinized and rehydrated. Antigen retrieval was performed at high heat (95°C) for 10 minutes in citrate buffer pH 6.0. Endogenous peroxidase activity was quenched with Bloxall (VectorLabs) for 10 minutes. Slides were blocked for 1 hour using 10% Normal Goat Serum (Invitrogen), incubated overnight with primary antibody. After 1-hour incubation with secondary antibody, VectaElite (VectorLabs) was applied on the sections for 30 minutes. Staining was visualized using DAB Peroxidase Substrate Kit (Vector Labs, SK-4100). Slides were counterstained with hematoxylin, dehydrated, and mounted with refrax mounting medium. Immunostained slides were scanned using a Perkin Elmer Slide Scanner (Panoramic MIDI Digital SlideScanner). Inform v2.1 image analysis software (Cambridge Research and Instrumentation) was used as a non-biased method to quantitate staining as previously described (*83*). Quantitation of c-MYC positive nuclei was performed using tumors of the same histological grade.

### RNA-sequencing

RNA from lung tumors and remaining lung tissues was isolated using Qiazol reagent using standard protocols (Qiagen cat # 799306). RNA purity was assessed by Agilent 2100 Bioanalyzer. Total RNA samples were sent to BGI Group, Beijing, China, for library preparation and sequencing. Reads (single-end 50bp at a sequencing depth of 20 million reads per sample) were generated by BGISEQ-500.

### RNA-seq analysis

Kallisto (https://pachterlab.github.io/kallisto/) was used to align to the reference transcriptome (ftp://ftp.ensembl.org/pub/current_fasta/mus_musculus/cdna/) and estimate transcript abundance. Differential gene expression analysis (DESeq2) was carried out using R (https://www.r-project.org/)). Differentially expressed genes were defined as having a adj. p-value<0.05 and fold change >+/-0.5. Gene ontology (GO) analysis was conducted on selected genes using the Database for Annotation, Visualization and Integrated Discovery (DAVID) (https://david.ncifcrf.gov/) program. Gene Set Enrichment Analysis (GSEA) (https://www.gsea-msigdb.org/gsea/index.jsp) was generated with TPM values from the above experiment using the java GSEA package.

### Lung nuclear extracts

Freshly collected lungs were mechanical homogenized in sucrose solution, and nuclei were isolated by ultracentrifugation through a denser layer of sucrose. Briefly, the whole lung was placed into a large (15 ml) dounce homogenizer on ice containing about 4 ml ice-cold PBS and 4 ml ice-cold homogenization solution (2.2 M sucrose with protease inhibitors, DTT and PMSF). Tissue was disrupted by pressing piston up/down 6x with loose piston and then 4x with tight piston. Homogenized tissue was added to an additional volume of ice-cold homogenization solution for a total volume of about 33 ml, which was then slowly poured on top of 10 ml cushion solution (2.05M sucrose with protease inhibitors, DTT and PMSF) in the ultracentrifugation tube (Beckman polyallomer #326823). After a 45-min spin at 24,600 rpm in pre-chilled SW32Ti rotor at 4°C, supernatants were carefully aspirated (the white pellets contain the nuclei). Nuclei were resuspended in 500 µl nuclear resuspension buffer (5mM Hepes pH 7.6; 50mM KCl and EDTA with DTT, protease inhibitors and PMSF) and transferred into a small (2 ml) dounce homogenizer on ice. Nuclei pellets were further resuspended by pressing piston up/down 3x with loose piston and then 2x with tight piston. Nuclei were then transferred to fresh 1.5 ml tubes and 500 µl of 2X NUN buffer (+ protease inhibitors and PMSF) were added while gently vortexing. After 20-minute incubation on ice, lysates were centrifuged for 20 minutes in ultracentrifuge at 38,000 rpm in pre-chilled 70.1Ti rotor with delrin adapters at 4°C. Supernatants were then transferred to clean tubes prior to protein quantification by BCA assay.

### Cell culture

A-427 and SK-LU-1 cells were purchased from the American Type Culture Collection (ATCC), cultured in Dulbecco’s modified Eagles medium (DMEM) plus 10% fetal bovine serum (Thermo Fisher) and 1% Pen-Strep (Gibco), and maintained in an atmosphere containing 5% CO_2_ at 37°C.

### Colony Formation Assay

Cells were plated into 6-well plates at 500 cells/well, and media was changed every 2 or 3 days with medium containing DTHIB (MedChemExpress, HY-138280) or corresponding concentration of DMSO. After indicated days of treatment, cells were washed in PBS, fixed for 10min with 100% methanol, and stained with 0.05% crystal violet for 20min. Plates were rinsed in DI-H_2_O, imaged and quantified using ChemiDoc XRS+ System (Bio-Rad).

### HSE-Luc activity

The pGL4.41[luc2P/HSE/Hygro] plasmid (HSE-Luc; Promega) was transfected into HEK293T cells and a stable clone expressing HSE-Luc was selected with hygromycin (200 mg/mL). For activity measurements, cells were plated in a flat, white, clear-bottom 96 well plate at a concentration of 50,000 cells/well. After 6 h, cells were pretreated with DTHIB (5 µM) and/or vehicle overnight. Compound A3 (10 µM; a kind gift from Rick Morimoto, Northwestern), MG132 (10 µM; Sigma), or DMSO was added for an additional 6 h. Plates were then equilibrated to room temperature and lysed by the addition of Bright-Glo reagent (100 µL; Promega) to each well. After a 10 min incubation to stabilize the signal, luminescence was then measured using an Inifinite F200 PRO plate reader (Tecan) and corrected for background signal.

### RNA extraction and quantitative RT-PCR

RNA was extracted from frozen tissues with Qiazol reagent using standard protocols (Qiagen cat # 799306). cDNA was prepared using QScript cDNA Supermix (VWR cat # 101414-106) and analyzed for gene expression using quantitative real-time PCR with iQ SYBR Green Supermix (Biorad cat # 1708885).

### Western Blots

Tissues or cells were lysed in RIPA buffer supplemented with protease and phosphatase inhibitors. Protein lysates were separated by SDS-PAGE and transferred to polyvinylidine difluoride (PVDF) membranes. Proteins were detected by standard Western blotting procedures. Antibodies were diluted 1:1,000 for BMAL1 (Abcam, ab93806), CRY1 and CRY2 (*84*), c-MYC (Abcam, ab32072), HSF1 (CST-12972), KRASG12D (CST-14429), Phospho-Erk1/2 (CST-4370), Erk1/2 (CST-4695); 1:2,000 for REV-ERBα (*33*), DNAJB1 (Enzo, ADI-SPA-400), HSP90AA1 (GTX109753); 1:10,000 for LAMIN A (Sigma, L1293); 1:50,000 for ACTIN (Sigma, A1978). Imaging and band quantification were carried out using ChemiDoc XRS+ System (Bio-Rad).

### Web-based analysis tools

Pathway analysis was performed with Enrichr (http://amp.pharm.mssm.edu/Enrichr). Quantifying circadian clock function using clock gene co-expression (CCD method) was carried using the web application available at https://hugheylab.shinyapps.io/deltaccd. Heatmap was generated by clustering using the Cluster 3.0 program (log2 transform data, center genes on mean, Hierarchical clustering with average linkage) (*85*), and then visualized with Java TreeView version 1.1.6r4 (*86*).

### Statistical analysis

Statistical analyses were performed using GraphPad Prism 8 software. Unless otherwise indicated, ANOVA was used to determine significance with a threshold of 0.05 acceptable false positive (P < 0.05). Rhythmicity was determined by JTK_Cycle analyses (*87*).

## Acknowledgments

We thank Drs. Lillian Eichner and Robert Svensson for helpful discussions and gifting us critical reagents and expertise. We thank Dr. Roderick Bronson for histopathological analysis of tumors, Nicole Madrazo for performing HSF1 luciferase assay, Toni Thomas, Judy Valecko, and Yolanda Slivers for administrative assistance, and Xia Jing for technical assistance.

## Author contributions

Conceptualization: MP, KAL

Methodology: MP, KAL, RJS, RLW, BJA

Investigation: MP, EI, RM, ABC, LHI

Visualization: MP, KAL, LHI, BJA

Supervision: KAL, RLW, RJS, MJB

Writing—original draft: MP, KAL

Writing—review & editing: MP, KAL, RJS, RLW, MJB, BJA

## Data and materials availability

RNA sequencing data are deposited in Gene Expression Omnibus (GEO) with accession number GSE194097.

