## Supplementary Materials for "Circadian disruption enhances HSF1 signaling and tumorigenesis in Kras-driven lung cancer"

**This PDF file includes:**

Figs. S1 to S9

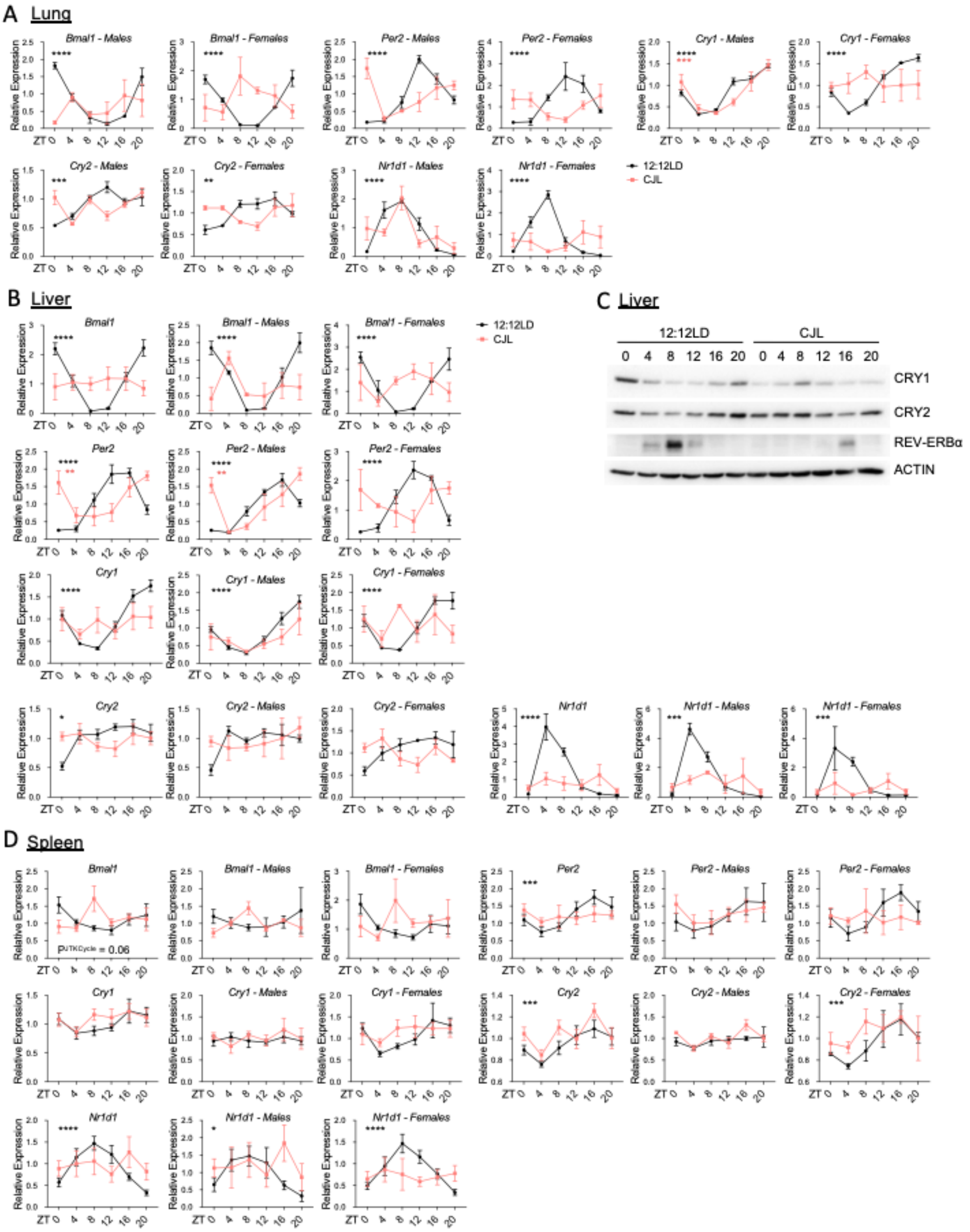


**Fig. S1. Core clock gene expression in tissues from mice housed in 12:12LD or chronic jetlag (CJL) conditions.**

C57BL/6J male and female mice were housed in 12:12LD or CJL for 8 weeks. Tissues were collected at the indicated zeitgeber times (ZT, hours after lights on) on Day 1 of the schedule shown in Fig. 1A. (A,B,D) Gene expression normalized to *U36b4* measured by quantitative realtime PCR from lungs (*A*), livers (*B*) and spleen (*D*). Data represent mean ± SEM for 3 males and 3 females per time point and light condition, both sexes plotted together or separately. Rhythmicity was determined by JTK_Cycle analyses; *$P^{JTKCycle} < 0.05$, **$P^{JTKCycle} < 0.01$, ***$P^{JTKCycle} < 0.001$ and ****$P^{JTKCycle} < 0.0001$. (C) Proteins detected by immunoblot from livers. Each lane on the Western blot represents a sample prepared from a unique animal. Representative images were taken from n = 6 biological replicates.

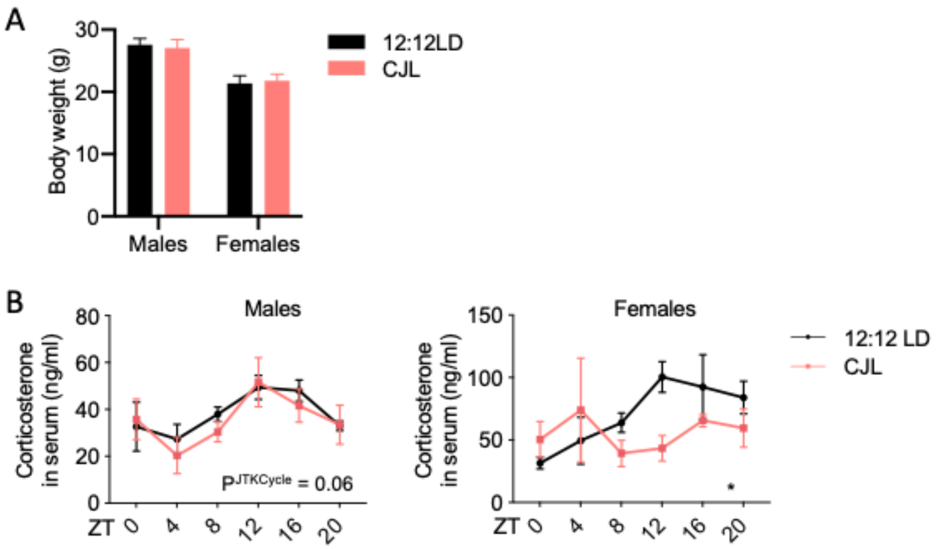


**Fig. S2. Body weight and serum corticosterone levels from mice housed in 12:12LD or chronic jetlag (CJL) conditions.**

C57BL/6J male and female mice were housed in 12:12LD or CJL for 8 weeks. **(A)** Weights were recorded prior to euthanasia. **(B)** Corticosterone levels in serum were measured from blood samples collected at the indicated times (hours after lights on) on Day 1 of the schedule shown in Fig. 1A. Data represent mean ± SEM for 3 males and 3 females per time point and light condition. Rhythmicity was determined by JTK_Cycle analyses; $*P^{JTKCycle} < 0.05$.

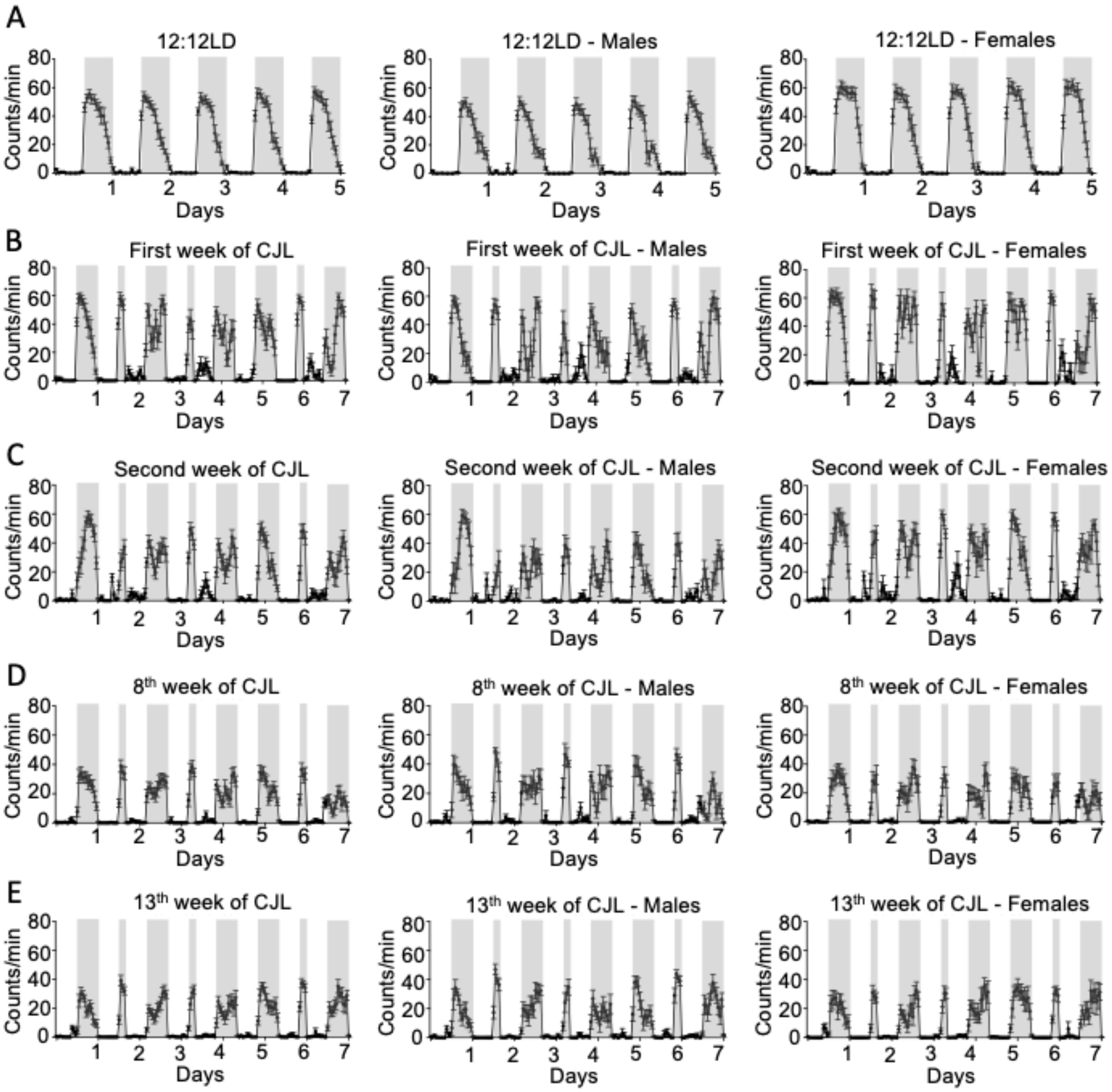


**Fig. S3. Wheel-running activity.**

(A-E) 8-weeks old C57BL/6J male and female mice were housed in 12:12LD for 2 weeks and then in CJL for 13 weeks in presence of a running wheel. Group mean waveforms of wheel running activity for the last 5 days in 12:12LD (A), first (B), second (C), eighth (D) and thirteenth (E) week in CJL conditions. Data represent mean ± SEM for 8 males and 8 females, plotted together or separately. Grey rectangles represent dark phases.

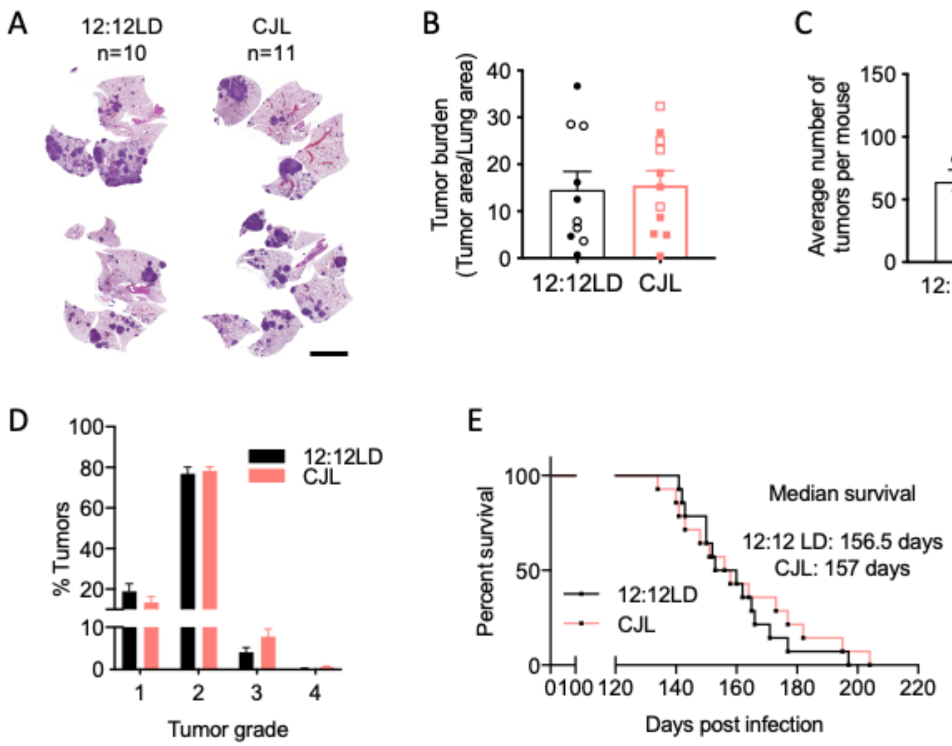


**Fig. S4. Tumor burden and overall survival in KP mice exposed to 12:12LD or CJL.**
Five weeks post-infection with lentivirus-Cre, KP mice were placed in either 12:12LD or CJL for 10 weeks (A-D) or until signs of distress (E). **(A)** Representative H&E-stained sections at endpoint; scale bar, 5000 µm. Tumor burden **(B)**, numbers **(C)**, and grade **(D)** were assessed from H&E sections. Column data represent mean ± SEM. Values for individual animals are plotted, clear and filled symbols represent males and females respectively. **(E)** Kaplan-Meier survival analysis for K mice placed in 12:12LD (n = 14) or CJL (n = 14) conditions.

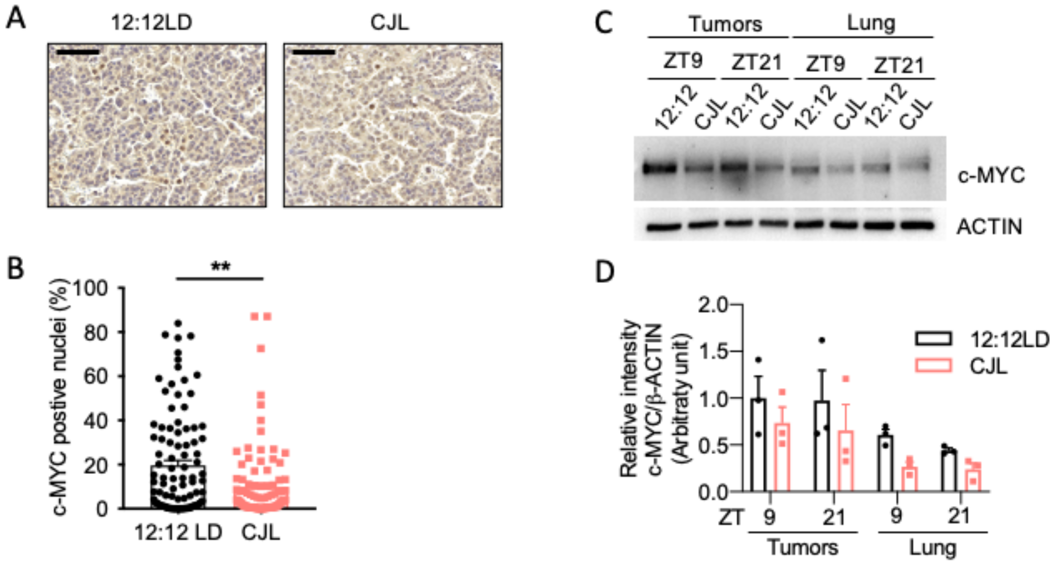


**Fig. S5. c-MYC protein levels in tumors and lungs from K mice exposed to CJL or normal light conditions.**

Five weeks post-infection with lentivirus-Cre, K mice were placed in either 12:12LD or CJL for 20 weeks. **(A)** Representative c-MYC IHC images in tumors from K mice sacrificed at ZT5; scale bar, 50 µm. **(B)** Quantitation of stained positive nuclei. **P <0.01 by Mann-Whitney test. **(C)** c-MYC detected by immunoblot. Tumors and lungs for each light condition and time point on the blot were from the same animal. Representative images were taken from n = 3 biological replicates. **(D)** Quantitation of (C).

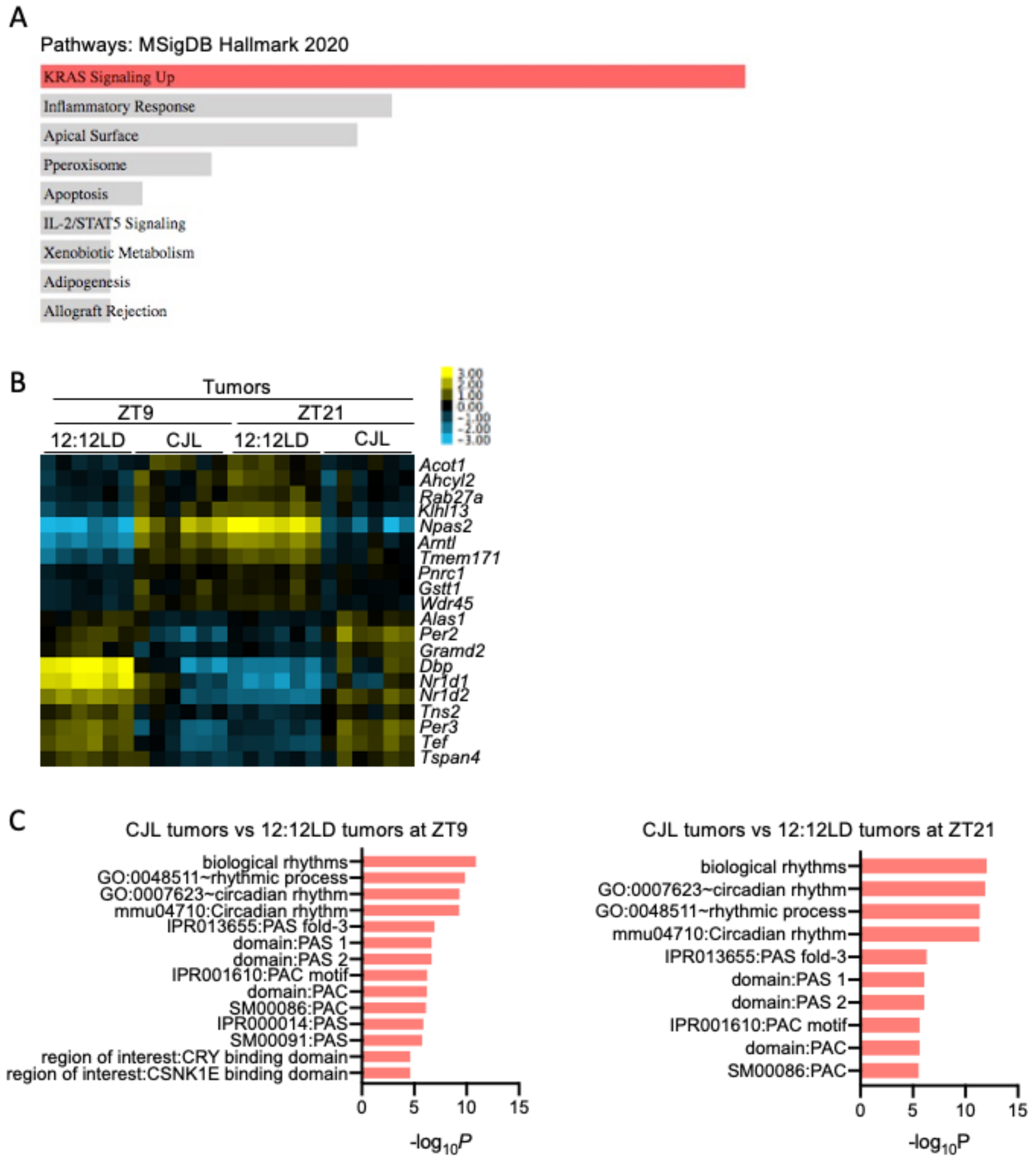


**Fig. S6. RNA-seq data analyses on lungs and tumors collected from K mice exposed to 12:12LD or CJL.**

Five weeks post-infection with lentivirus-Cre, K mice were placed in either 12:12LD or CJL for 20 weeks. (A) Enrichr Pathway analysis of the upregulated genes in tumors compared to whole lungs (48 genes with log2FoldChange >= 1.5), all conditions combined, sorted by p-value ranking. Colored bar correspond to terms with significant p-values (<0.05). (B) Heatmap of the 20 genes differentially expressed upon CJL versus 12:12LD in tumors at both ZT9 and ZT20. **(C)** DAVID analyses on the differentially expressed genes in tumors between 12:12LD and CJL. Only terms with FDR< 0.25 are shown.

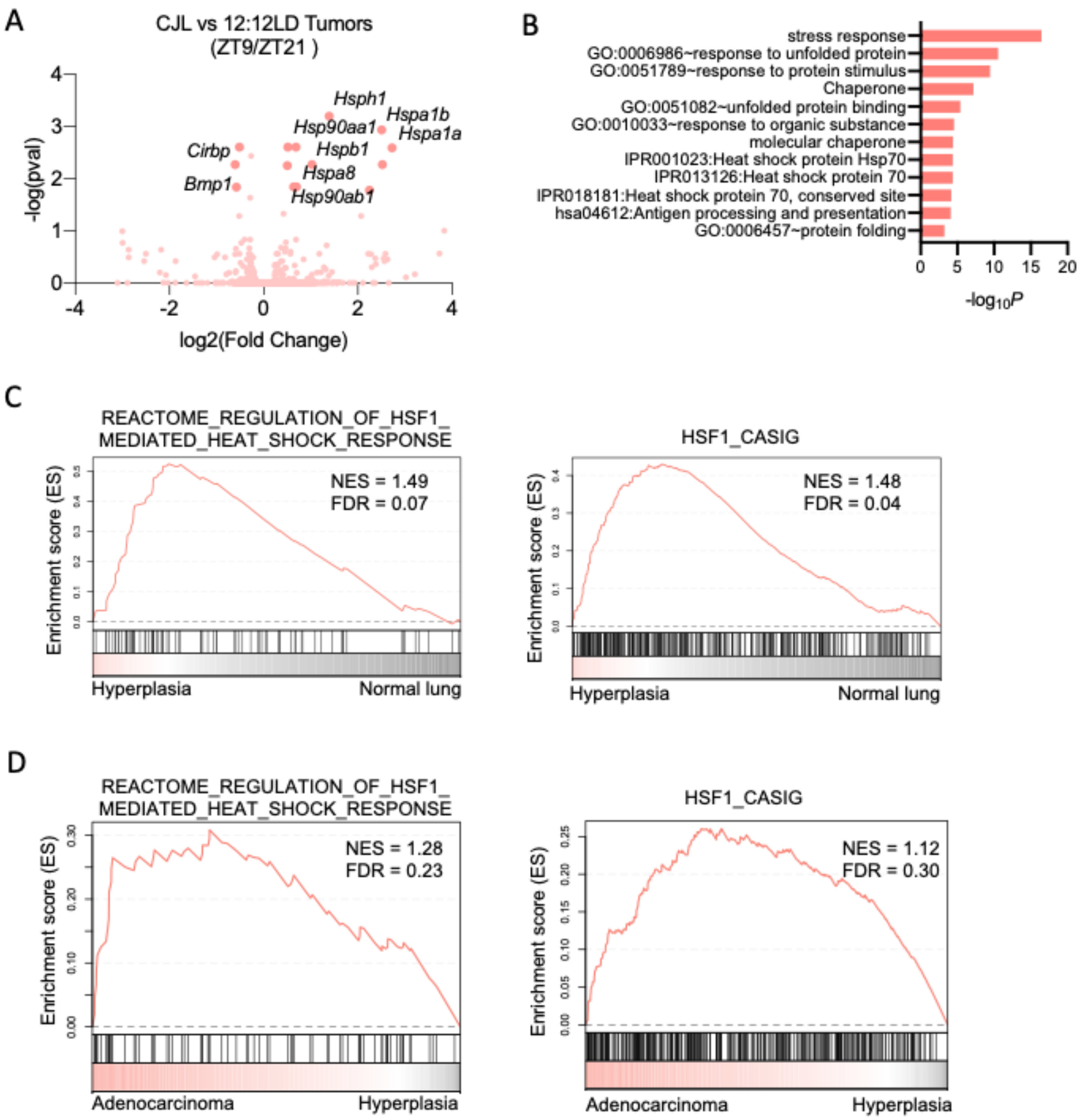


**Fig. S7. Up-regulation of HSF1-mediated heat shock response and cancer signature in mouse models of Kras-driven lung cancer.**

**(A,B)** Five weeks post-infection with lentivirus-Cre, K mice were placed in either 12:12LD or CJL for 20 weeks. **(A)** Volcano plots of differentially expressed genes between 12:12LD and CJL by DESeq2 analyses for tumors only, taking time of collection (ZT9/21) as confounding factor. **(B)** DAVID analyses on the differentially expressed genes by DESeq2 in tumors between 12:12LD and CJL, taking time of collection as confounding factor. Only terms with FDR< 0.25 are shown. **(C-D)** GSEA plots for the HSF1-mediated heat shock response reactome and HSF1-Cancer signature gene sets applied to Ambrogio et al., 2016 microarray data of laser-capture microdissected hyperplasic or normal lung cells (C) and adenocarcinoma or hyperplasic lesions (D) from K-Ras$^{G12V}$ mice..

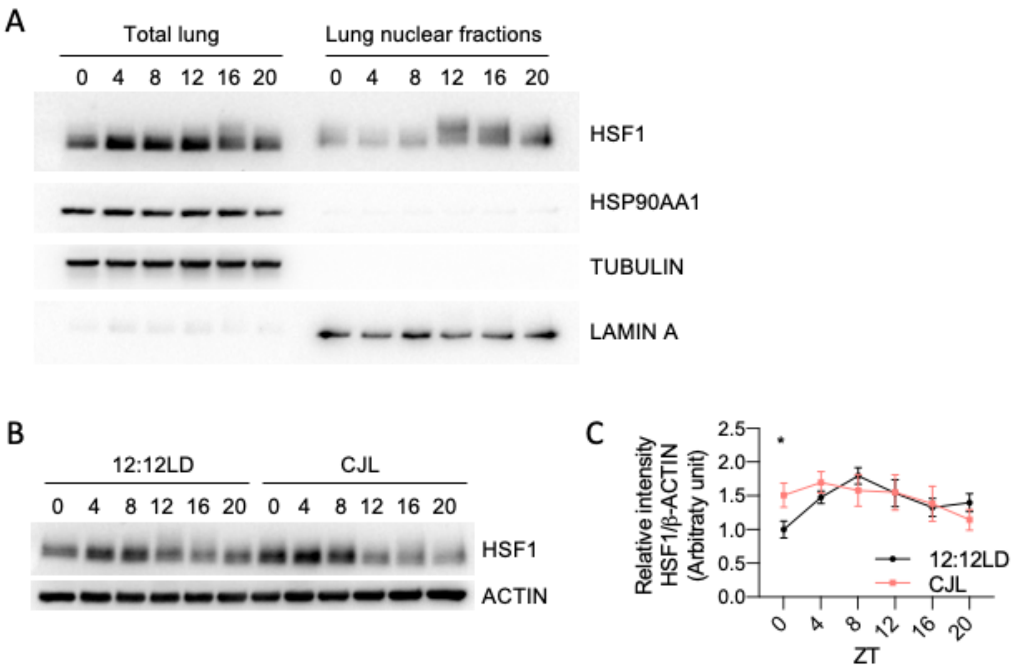


**Fig. S8. HSF1 protein levels in lung.**

**(A)** Proteins from whole lung or lung nuclear extracts detected by immunoblot from C57BL/6J mice housed in 12:12LD for 8 weeks. Lung tissues were collected at the indicated times (hours after lights on). Each lane on the Western blot represents a sample prepared from a unique animal. **(B)** Proteins from whole lungs detected by immunoblot from the same cohort of C57BL/6J as in Fig. 1. Each lane on the Western blot represents a sample prepared from a unique animal. Representative images were taken from *n* = 3 biological replicates. **(C)** Quantitation of (B). Rhythmicity was determined by JTK_Cycle analyses; $*P^{JTKCycle} < 0.05$.

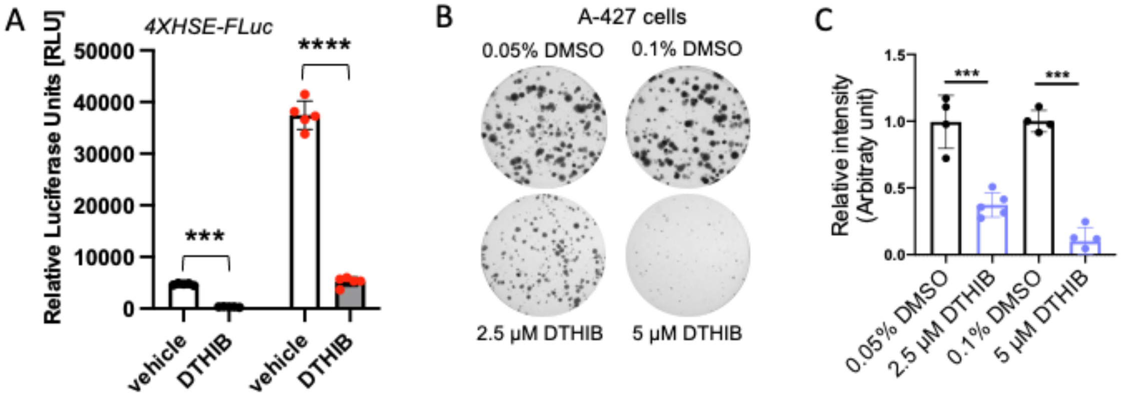


**Fig. S9. Pharmacological inhibition of HSF1 by DTHIB.**

**(A)** Relative luciferase activity in HEK293 cells stably expressing the reporter *pGL4.41Luc2P/4XHSE/Hygro* and treated with vehicle (black symbols) or the HSF1 activator A3 (red symbols) and vehicle (open bars) or 5 µM DTHIB (filled gray bars). *** $P < 0.001$, **** $P < 0.0001$ by two-way ANOVA. **(B)** Representative images of crystal violet-stained colonies formed by A-427 cells treated with DTHIB or vehicle DMSO 7 days after seeding, for 9 days. **(C)** Quantification of (B) from three biological replicates. Each condition was compared to controls that were plated in wells on the same plates. Bars represent mean ± SD, **$P < 0.01$ by student t-test.
